# Innate immunity can distinguish beneficial from pathogenic rhizosphere microbiota

**DOI:** 10.1101/2023.01.07.523123

**Authors:** David Thoms, Melissa Y. Chen, Yang Liu, Zayda Morales Moreira, Youqing Luo, Siyu Song, Nicole R. Wang, Cara H. Haney

## Abstract

For optimal growth and development, hosts depend on their ability to promote healthy symbiotic interactions while restricting pathogen growth. To ask whether hosts can distinguish phylogenetically similar pathogens and beneficial bacteria, we used two closely related plant-associated strains of *Pseudomonas fluorescens* where one is beneficial and the other exhibits toxin-dependent virulence. We show that while the two strains co-exist *in vitro*, the beneficial outcompetes that pathogen *in planta*. Using several readouts for plant innate immunity, we found that the beneficial and pathogenic strains elicit mechanistically distinct immune responses that occur in distinct root compartments. We show that while both the pathogenic and beneficial bacterial have plant recognizable MAMPs, the pathogen uniquely induces MAMP-independent immune responses. We found that the pathogen induces both a toxin-independent and a unique toxin-dependent defense response that remains intact in immune signaling mutants including *bak1/bkk1* and *npr1/4D*. We conclude that hosts can distinguish between phylogenetically similar microbes.

## Introduction

Hosts rely on a healthy microbiome to facilitate growth, development, and fitness. Plants and animals rely on functionally analogous innate immunity systems to distinguish self from non-self ^1,2^. The animal gut and plant root are important sites of nutrient acquisition that are colonized by dense microbial communities that may aid or hinder the host. Recent work suggests that the innate immune system plays a key role in controlling both host-microbe and microbe-microbe interactions in a manner that promotes beneficial interactions and limits disease ^3–8^. However, how the innate immune system distinguishes between helpful and harmful microbes to establish a beneficial microbiome is poorly understood. As plants lack an adaptive immune system, they are useful models for isolating the role of innate immunity in shaping the host microbiome.

It is not entirely clear which innate immune mechanisms would be most important for distinguishing beneficial microbiota from pathogens. Plants sense potential pathogens through intracellular and extracellular receptors that activate signaling pathways commonly referred to as effector-triggered immunity (ETI) and pattern-triggered immunity (PTI), respectively ^9,10^. Effectors are microbe-derived molecules that alter host physiology in a manner that benefits the pathogen. ETI occurs via direct or indirect effector recognition, which allows plants to detect highly specialized pathogens; however, many opportunistic bacterial pathogens use effector-independent virulence mechanisms ^11–13^. PTI is based on receptor-mediated recognition of evolutionarily conserved microbe-associated molecular patterns (MAMPs). Yet, as MAMPs are highly conserved between pathogens and commensals, they provide limited resolution for distinguishing between harmful and beneficial microbes. Furthermore, the horizontal transfer of non-effector virulence genes ^14^ can lead to pathogens that share MAMPs with closely related non-pathogenic strains. This loss and gain of toxins via horizontal gene transfer events is ubiquitous in nature ^15–18^, and presents a challenge for innate immunity to distinguish between harmful and beneficial lifestyles through MAMPs and effectors. As a result, it is currently unclear how PTI and ETI could allow for detecting and restricting pathogens in the presence of complex, quickly evolving microbiota. It is likely that immune mechanisms other than PTI and ETI are used to recognize harmful microbes within diverse microbial communities.

In this work, we use a system in which the gain and loss of genomic islands confers either pathogenic or beneficial lifestyles among two closely related strains of *Pseudomonas fluorescens* ^15^. The pathogen, *P. fluorescens* N2C3, lacks a type III secretion system and virulence was acquired via quorum-sensing regulated expression of lipopeptide toxins, syringomycin and syringopeptin. The beneficial, *P. fluorescens* WCS365, lacks the genomic islands that confirm pathogenicity, but has genetic traits associated with commensalism. When co-inoculated with pathogenic *P. fluorescens* N2C3, the beneficial WCS365 protects plants from N2C3 pathogenesis in a colonization dependent manner, suggesting that the protection is driven by competition between the beneficial and pathogen ^19^. As no evidence was found for direct microbial antagonism between these strains ^19^, we hypothesized that outcome of competition may in part be due to a plant-dependent mechanism rather than microbe-microbe interactions. Here we show that plants are able to distinguish between pathogenic and beneficial bacteria via both toxin-dependent and independent mechanisms. Our data indicate that plants can distinguish closely related beneficial and pathogenic bacteria independently of MAMP or effector recognition. We propose that the presence of bacterial toxins or toxin-mediated damage may allow plants to broadly distinguish between pathogenic and beneficial microbiota.

## Results

### Beneficial *Pseudomonas* outcompeting a pathogen is plant dependent

To determine how hosts can distinguish between bacterial lifestyles acquired over short evolutionary time scales, we use the model plant *Arabidopsis thaliana*, and two closely related *Pseudomonas* strains: beneficial *P. fluorescens* WCS365 and pathogenic *P. fluorescens* N2C3. When co-inoculated onto plant roots, WCS365 protects plants from N2C3 pathogenesis ^19^. To test whether protection is plant-dependent, we used coculture competitions *in vitro* and *in planta*. To quantify bacterial growth, *P. fluorescens* WCS365 was transformed with a plasmid with or without the mTurquoise2 fluorescent reporter, and then competed against other bacterial strains expressing an empty vector. Fluorescence-based calculations of optical density (OD) of WCS365 were used for growth comparisons.

In agreement with prior findings ^19^, we found that *P. fluorescens* WCS365 significantly outcompeted N2C3, but not itself, *in planta* (Figure 1A, B). *In planta P. fluorescens* WCS365 reached stationary phase in ~20% less time when competed against N2C3 than in competitions against itself or when grown in monoculture (Figure 1A). By day five, the final OD_600_ of WCS365 was significantly higher when cocultured with N2C3 compared to competitions against itself (Figure 1B; p < 0.002). The area under the curve (AUC) for WCS365 growth in the presence of N2C3 (AUC: 1.08, CI: 0.41 – 1.75) was ~50% higher compared to competition against itself (AUC: 0.70 CI: 0.47 – 0.93) (Figure 1A). These data indicate that by quantifying total bacterial growth, beneficial *P. fluorescens* WCS365 robustly outcompetes pathogenic N2C3 *in planta*.

**Figure 1.**
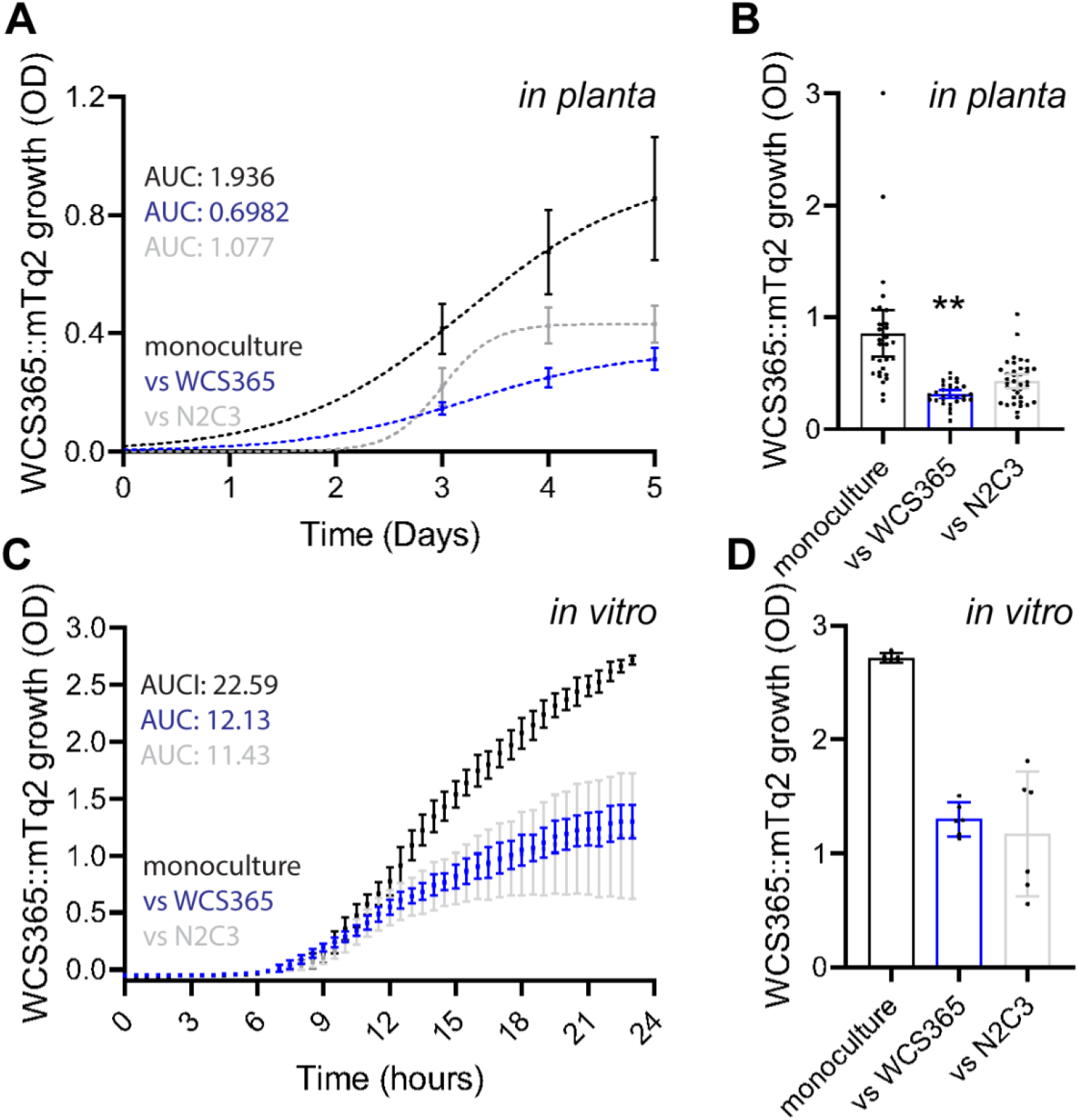
Beneficial *P. fluorescens* WCS365 outcompetes pathogenic N2C3 *in planta* but not *in vitro*. Fluorescence-based competition was measured between a wildtype WCS365 strain containing an empty expression vector and wildtype WCS365, wildtype N2C3, or N2C3 ΔSYRΔSYP mutant strains containing a plasmid expressing mTurquoise2. A and B, Growth curves of 5-day *in planta* competitions fitted with a logistic growth model. Data points are of individual biological replicates, from two temporal replicates. C and D Growth curves of 23-hour *in vitro* competitions. Data points are of means (C) or individual biological replicates (D) pooled from five experimental replicates. A and C, Monoculture WCS365 shown in black; competitions between WCS365 versus self in gray; WCS365 versus N2C3 in blue. Area under the curve (AUC) are shown for each bacterial growth curve. B and D, WCS365 final OD at 5 DPI for monoculture and coculture experiments. A Welch’s t test was used to determine statistical significance, ** *p* < 0.01. All error bars show 95% confidence intervals.

In contrast, we found that the beneficial *P. fluorescens* WCS365 did not outcompete N2C3 in 23-hour *in vitro* competitions. WCS365 growth was similar when cocultured with itself (AUC: 12.13, 95% CI: 11.74 – 12.52) compared to when cocultured with N2C3 (AUC: 11.43, 95% CI: 10.28 – 12.59) (Fig 1C, D). There was also no significant difference (*p* = 0.59) for the final OD_600_ of WCS365 at the end of the 23-hour assay when cocultured with N2C3 compared to competitions against itself. Therefore, while WCS365 does not outcompete N2C3 *in vitro*, it does outcompete *in planta* indicating that the competition is dependent on plant immunity or a plant-specific nutrient.

### Pathogenic *P. fluorescens* N2C3 can overcome spatial restrictions in rhizosphere immunity

Prior work demonstrates a clear compartmentalization of immunity in the animal gut and plant root ^20,21^. While defense-associated responses to bacterial MAMPs are typically restricted to the root elongation zone ^22,23^, the root maturation zone can also exhibit an immune response when wounding and MAMPs co-occur ^24^. We used the MAMP-inducible fluorescent defense reporter line *pPER5::mVenusx3* to observe immunity compartmentalization in the root in response to *P. fluorescens* to determine whether beneficial WCS365 or pathogenic N2C3 induces a defense response. We used the plant-derived damage-associated molecular pattern (DAMP) AtPep1, as a positive control which induces a potent immune response throughout the root ^22^. Through quantifying the fluorescence signal along the root (Figure 2A) and calculating total fluorescence by measuring the area under the curve (Figure 2B), we found that pathogenic *P. fluorescens* N2C3 and the AtPep1 strongly induced the *PER5* promoter in the maturation zone, while beneficial *P. fluorescens* WCS365 and mock treatment did not (Figure 2C). WCS365, N2C3, and AtPep1 all significantly induced the *pPER5* in the root tip compared to mock treatment (Figure 2C). This suggests that PER5 induction by pathogenic *P. fluorescens* N2C3, like AtPep1, overcomes immune compartmentalization in the root. These data indicate that a differential defense response to WCS365 and N2C3 is spatially regulated and suggests that N2C3 virulence can overcome spatially restricted immune barriers.

**Figure 2.**
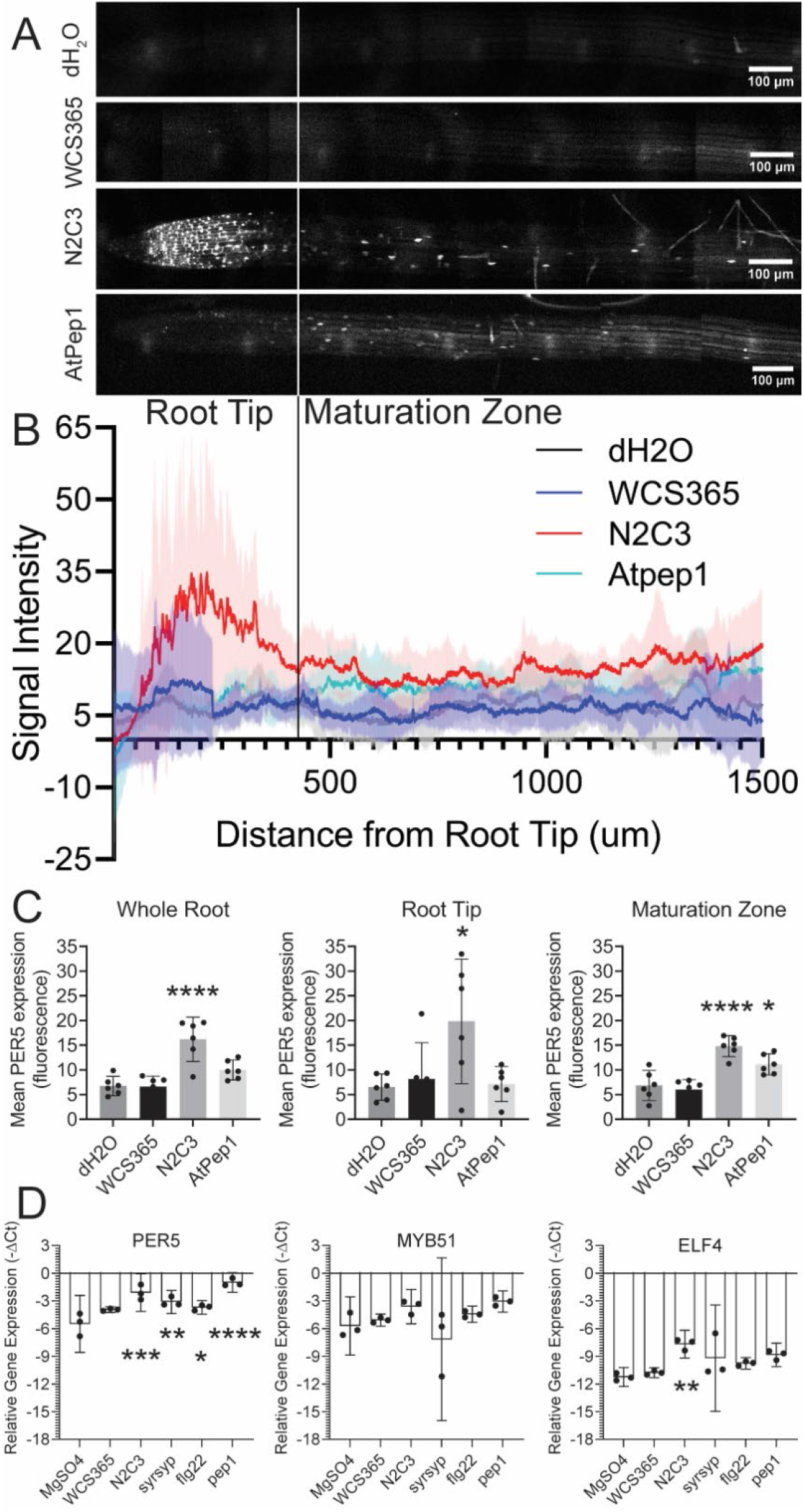
N2C3 and WCS365 induce distinct spatial expression profiles of defense gene expression in roots. A, fluorescence images of *pPER5::mVenusx3* nuclear-localized expression in root cells two days after bacteria inoculation and 4.5 hours after AtPep1 treatments. B, Fluorescence-based quantification of *pPER5::mVenusx3* expression (fluorescence) along the root ranging from the root apical meristem to the maturation zone. C, Area under the curve (AUC) measurements of *pPER5::mVenusx3* fluorescence data shown in B, separated by root compartments. D, RT-qPCR data of defense-associated relative gene expression (-ΔCt) in the root maturation zone comparing bacteria and elicitor treatments in wildtype plants. Increased -ΔCt values show increased gene expression. Statistical significance was determined with a one-way ANOVA comparing mean fluorescence intensity or -ΔCt of bacterial or elicitor treatments to mock, \**p* < 0.05, \*\**p* < 0.01, \*\*\**p* < 0.001, \*\*\*\**p* < 0.0001. All error bars show 95% confidence intervals.

To determine if *P. fluorescens* N2C3 also induces expression of other defense-associated genes in the root maturation zone in a syringomycin and syringopeptin-dependent manner, we used RT-qPCR to quantify expression of three defense associated genes (*PER5, MYB51, ELF4*). In agreement with the fluorescent reporter line, beneficial WCS365 did not induce expression of *PER5, MYB51*, and *ELF4* expression (*p* > 0.05) (Figure 2), while pathogenic N2C3 infection induced expression of *PER5* (*p* < 0.001) and *ELF4* (*p* < 0.005). The N2C3 Δ*syr*Δ*syp* avirulent mutant (*p* < 0.01) and flg22 (*p* < 0.05) treatments significantly induced expression of *PER5*. The Δ*syr*Δ*syp* mutant has a deletion of syringomycin and syringopeptin resulting in a loss of virulence ^15^. AtPep1 treatment significantly induced PER5 (*p* < 0.0001) and showed an increased trend for *ELF4* (*p* = 0.052). Collectively, these results indicate that the defense response to N2C3 is compartmentalized in the root and triggered by syringomycin- and syringopeptin-mediated virulence.

### Pathogenic N2C3 but not beneficial WCS365 induces a damage-like response in the root

Our data show that the root maturation zone exhibits a differential response to beneficial *P. fluorescens* WCS365 and pathogenic N2C3 in a syringomycin- and syringopeptin-dependent manner. Syringomycin and syringopeptin are pore-forming toxins that can disrupt eukaryotic cells by oligomerizing into plasma membrane channels ^25–27^. To determine if the N2C3-induced defense response resembles a PTI or damage-like response, we used RNAseq to quantify gene expression after treatment with bacterial strains and immune elicitors. We used flg22 as an elicitor for PTI and AtPep1 as an elicitor for damage-triggered immunity. Overall, we found that roots treated with the pathogen had the largest transcriptomic response (10,038 Differentially Expressed Genes [DEGs]) (Figure 3A). Roots treated with WCS365 (93 DEGs) or flg22 (4 DEGs) induced little change in plant gene expression in the maturation zone relative to the mock treated control. In agreement with prior work, AtPep1 induced a large number of genes (3018 DEGs) in the root maturation zone (Figure 3A).

**Figure 3.**
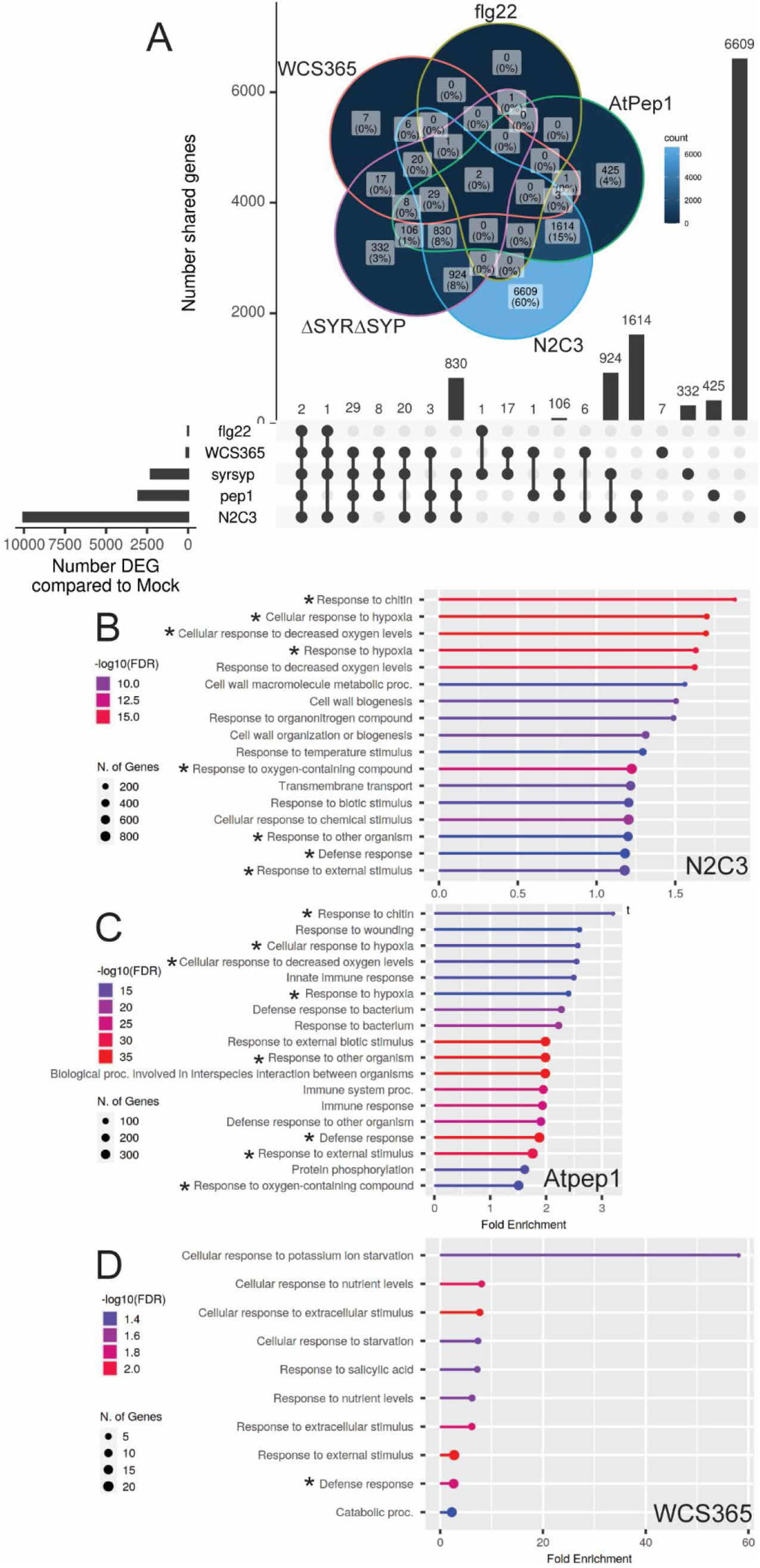
The *P. fluorescens* pathogen, but not the beneficial, induces a strong damage-like response in roots. A, Upset plot and Venn Diagram showing the number of differentially expressed genes (DEGs) in the root maturation zone comparing treatments with WCS365, N2C3, Δ*syr*Δ*syp* N2C3, flg22, or AtPep1, to mock. B – D, ShinyGo analysis of all significant DEGs from roots treated with N2C3 (B), AtPep1 (C), or WCS365 (D) showing biological processes GO term enrichments. Significant DEGs were selected based on the DESeq2 adjusted *p* value (*p* < 0.01). GO terms that are shared between AtPep1 and bacterial treatments are marked with an asterisk.

Relative to mock-treated roots, we found that genes induced by pathogenic *P. fluorescens* N2C3 treatment had a significant overlap with a damage elicitor and induced 82.1% (2478/3018 DEGs) of the genes induced by AtPep1 treatment and 75.0% (3/4 DEGs) of the genes induced by flg22 treatment (Figure 3A). Upregulated and downregulated DEGs followed a similar trend, with 79.6% (1533/1926 DEGs) of upregulated AtPep1 genes and 84.2% (865/1027) of downregulated AtPep1 genes also induced by N2C3 treatment (Supplemental Figure 1A). Of the plant DEGs induced in response to the avirulent N2C3 Δ*syr*Δ*syp* mutant, 18.0% (1806/10,038 DEGs) were shared with wildtype N2C3, and 32.3% (975/3018 DEGs) were shared with AtPep1. These results show a strong overlap between the N2C3 pathogen and AtPep1 induced genes occurring in a syringomycin- and syringopeptin-dependent manner.

Since more than three quarters of genes induced by virulent *P. fluorescens* N2C3 treatment were not induced by AtPep1 treatment, we hypothesized that N2C3 might also induce a set of biological processes distinct from AtPep1 treatment. To test this, we used the ShinyGO ^28^ gene ontology (GO) pathway enrichment analysis to compare biological processes GO terms between the two treatments. We found that N2C3 and AtPep1 induced very similar biological processes (e.g. response to: chitin, hypoxia, biotic stimulus, etc.), with few processes unique to virulent N2C3 (e.g. cell wall biogenesis, temperature stimulus, transmembrane transport) or AtPep1 (e.g. wounding response, bacterium response, protein phosphorylation) treatment (Figure 3B, C). Therefore, despite the difference in magnitude of the transcriptomic responses to the pathogen and AtPep1, the two treatments induced very similar biological processes suggesting pathogenic N2C3 induces a damage-like immune response.

In contrast, the root maturation zone showed a very small transcriptomic response to the beneficial *P. fluorescens* WCS365 (93 DEGs). Plant treatment with WCS365 only induced 1.4% (43/3018 DEGs) of AtPep1 DEGs and 75% (3/4 DEGs) of flg22 induced genes (Figure 3A). Similarly, 1.1% (22/1926 DEGs) of upregulated AtPep1 genes and 1.5% (15/1027) of downregulated AtPep1 genes were induced by WCS365 treatment (Figure 3C, D). Interestingly, 62.4% (58/93 DEGs) of WCS365-induced genes were also induced by the virulent and avirulent pathogen strains. Over half (31/58 DEGs) of these *P. fluorescens* inducible genes were also induced by AtPep1. These results suggest that the majority of beneficial *P. fluorescens* WCS365 DEGs are generally induced by both beneficial and pathogenic *P. fluorescens*.

Since beneficial *P. fluorescens* WCS365 induced a relatively small transcriptomic response we hypothesized that WCS365 would not induce similar biological processes as N2C3 or AtPep1. In agreement with our hypothesis, ShinyGO analysis showed several unique biological processes (e.g. response to: potassium starvation, nutrient levels, salicylic acid, etc.) induced by only WCS365 (Figure 3D) but not by N2C3 or AtPep1 treatments (Figure 3B, C). However, beneficial *P. fluorescens* WCS365 also induced a small set of genes associated with a defense response, consistent with our finding that 46.2% of WCS365 DEGs were AtPep1 inducible (Figure 3A). Gene set enrichment analysis showed a weak overlap in biological response to WCS365 when compared to N2C3 and AtPep1 treatments (Supplemental Figure 1B). Cumulatively, these results show that while roots induce a small subset of defense-associated genes in response to *P. fluorescens*, the pathogen, but not the beneficial strain, induces a potent transcriptomic response that induces a set of biological processes similar to DAMP-triggered immunity.

### Plants respond to pathogenic *P. fluorescens* N2C3 in a toxin- dependent and - independent manner

Since pathogenic N2C3 treatment strongly induced expression of AtPep1-associated genes (Figure 3A), we hypothesized that the core transcriptomic response to N2C3 would be an immune response. However, since N2C3 treatment resulted in such a large transcriptomic response in the RNAseq dataset, we used a principal component analysis (PCA) to reduce the number of variables representing the data. The principal component 1 axis (PC1) explained 61% of the variance in the dataset, while the PC2 axis explained 10% of the variance. Surprisingly, the virulent and avirulent N2C3 treatments showed distinct separation in the PCA, with the transcriptomic response to the virulent strain distinct from all other treatments along PC1 and the response to the avirulent strain clustering with AtPep1 treatment along PC2 (Figure 4A). This separation in responses to virulent and avirulent N2C3 strains suggests that roots exhibit two separate transcriptomic responses to the pathogen in a manner dependent on the presence or absence of syringomycin and syringopeptin toxins. Therefore, we hypothesized that the pathogen would induce both toxin-dependent and -independent immune responses.

**Figure 4.**
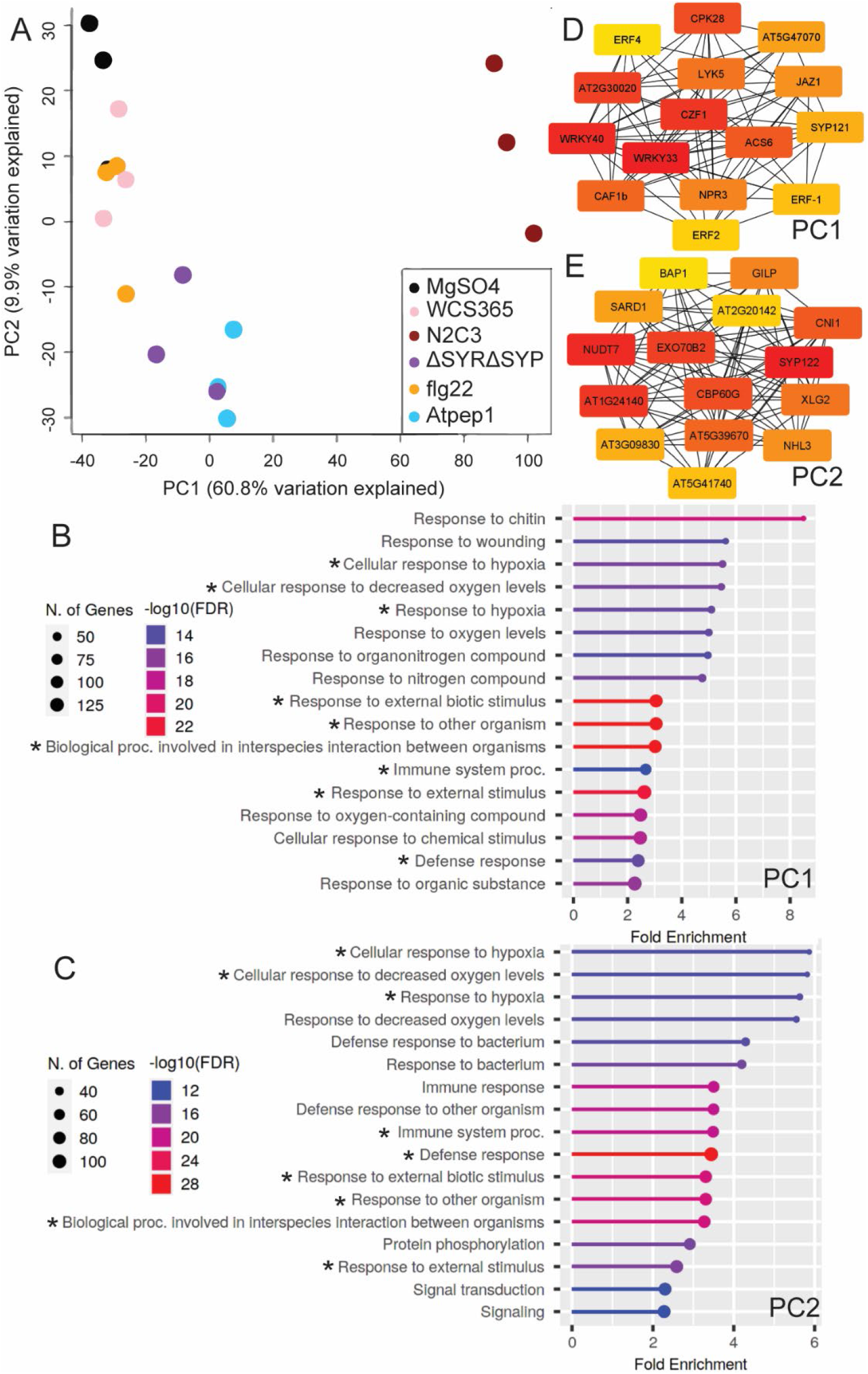
Pathogenic *P. fluorescens* N2C3 induces root immunity via syringomycin- and syringopeptin-dependent and -independent mechanisms. A, Principal component analysis for all significantly induced DEGs (Figure 3) from roots treated with mock, WCS365, N2C3, Δ*syr*Δ*syp* N2C3, flg22, or AtPep1. B and C, ShinyGO enrichment analysis of wildtype N2C3 DEGs strongly associated with PC1 (B) or AtPep1 and Δ*syr*Δ*syp* N2C3 induced genes associated with PC2 (C). D and E, Node network analysis of the 15 highest scoring nodes from PC1 (D) or PC2 (E) associated genes. Red nodes = high scores, yellow = lower scores. FDR = false discovery rate. Shared GO terms are marked with an asterisk.

To determine if the transcriptomic response to virulent N2C3 encompassed an immune or susceptibility response, we identified all genes whose highest association was with PC1 and cross-referenced these genes with our DEseq2 list of significantly induced DEGs. This produced a shortlist of 740 significantly induced DEGs of interest, all of which were either uniquely upregulated by wildtype N2C3 or by both AtPep1 and wildtype N2C3 treatments (Supplemental Table 1). We will refer to the genes derived from the PC1 list as syringomycin- and syringopeptin-dependent genes. Likewise, we used the PC2 axis to determine if the transcriptomic response to avirulent N2C3 was associated with immunity. We selected genes negatively associated with PC2 if they were induced by AtPep1 or the Δ*syr*Δ*syp* mutant treatment, since the Δ*syr*Δ*syp* mutant and AtPep1 treatments clustered along the PC2 axis. This produced a unique list of 548 genes (Supplemental Table 2), which we will refer to as syringomycin- and syringopeptin-independent genes. We used ShinyGO enrichment analysis of the syringomycin- and syringopeptin-dependent and independent gene lists to identify the presence or absence of biological processes associated with immunity. In agreement with our hypothesis, both the syringomycin- and syringopeptin-dependent (Figure 4B) and independent (Figure 4C) responses showed strong enrichment for terms associated with immunity (e.g. defense, oxidative stress, interspecies interaction, etc.). Gene set enrichment analysis showed a strong overlap between the syringomycin- and syringopeptin-dependent and independent biological response (Supplemental Figure 1C). These data suggest that plants respond to the N2C3 pathogen in a syringomycin and syringopeptin-dependent and -independent manner.

To determine if the transcriptomic response unique to N2C3 was due to the host being in a susceptible state, we cross referenced the syringomycin and syringopeptin dependent and independent gene lists with immunity and susceptibility transcriptional responses in leaves ^29^. We found that the syringomycin- and syringopeptin-dependent response induced 10/133 genes associated with immunity and 1/28 gene associated with susceptibility (Supplemental Table 3). The syringomycin- and syringopeptin-independent response induced a different set of 15/133 genes associated with immunity and 0/28 genes associated with susceptibility (Supplemental Table 4). This lack of overlap suggests that the plant response to the pathogen does not resemble susceptibility in leaves. Taken together, these data suggest that N2C3 induces a unique syringomycin- and syringopeptin-dependent defense response that is distinct from the less potent syringomycin- and syringopeptin-independent defense response.

Despite induction of a defense response by both virulent and avirulent strains, our data suggest that the presence or absence of virulence leads to distinct transcriptomic responses (Figure 3A, 5A). To determine if the presence or absence of syringomycin and syringopeptin were inducing a similar defense response, we identified hub genes as potential key regulators for the syringomycin and syringopeptin -dependent and - independent responses. We found WRKY33 and WRKY40 to have the highest degree score (network connections) for syringomycin and syringopeptin dependent defense response (Figure 4D). Node analysis of the syringomycin- and syringopeptin-independent defense response identified several genes important for regulating salicylic acid (SA) signaling and the hypersensitive response (Figure 4E). There was no overlap among the top 15 hubs genes in the syringomycin- and syringopeptin-dependent and - independent gene lists (Figure 4D, E). These results suggest that N2C3 induces two distinct defense responses dependent on the presence or absence of syringomycin and syringopeptin.

**Figure 5.**
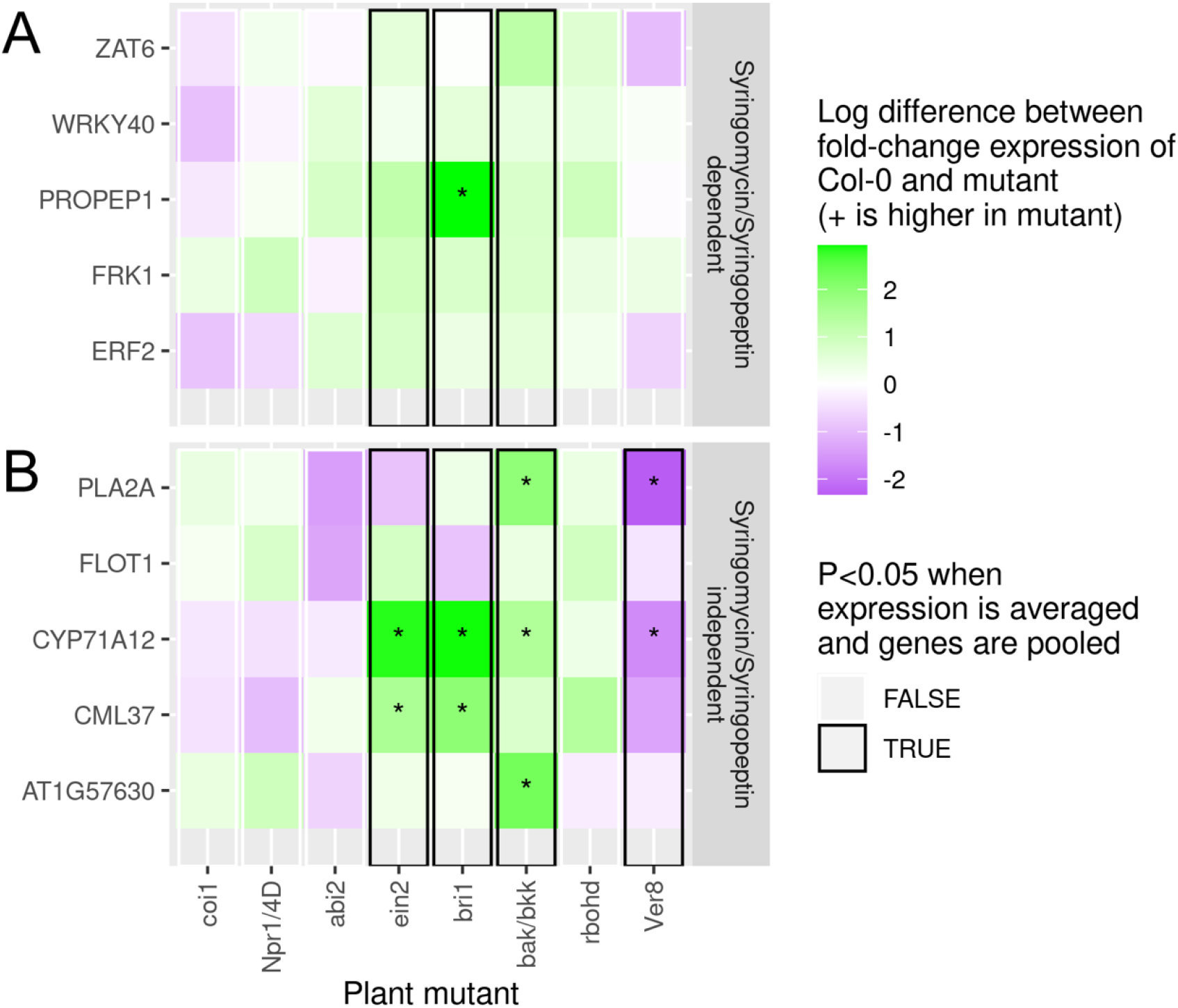
Syringomycin- and syringopeptin-dependent response to pathogenic N2C3 is inhibited by BAK1/BKK1 and ethylene. A, B, RT-qPCR heatmaps of the root maturation zone response to N2C3 across several plant genotypes, two days post treatment. Reporter genes for the response to N2C3 were selected via the PCA of the RNAseq data (Fig. 4D). Five genes were selected from as reporters for the syringomycin- and syringopeptin-dependent immune response to N2C3, while five other genes were selected from as reporters for syringomycin- and syringopeptin-independent immune response to N2C3. Colormap show the difference of relative log2 fold change (ΔΔΔCt ^62,63^) calculated from the difference in gene expression between N2C3 and MgSO_4_ treatments. A positive ΔΔΔCt value indicates an increase in gene expression in the mutant line. Linear regression models in the R programming language were used to determine significance between Col-0 and each plant mutant line comparing either the single or the collective five-gene means (Significant individual genes indicated by \**p* < 0.05, with significant pooled genes boxed in black).

### The defense response to N2C3 is not dependent on PTI or hormone signaling

To identify plant pathways required for the defense response induced by pathogenic *P. fluorescens* N2C3, we tested N2C3 toxin dependent- and independent-gene expression in plant mutants defective in key hormone pathways [*coi1-16* (JA-insensitive) ^30^, *npr1/4D* (SA-insensitive) ^31^, *abi2* (ABA-insensitive) ^32^, *ein2* (ET-insensitive) ^33^, *bri1* (BR-insensitive) ^34^] and immunity associated components [*bak1-5/bkk1* (PTI deficient)^35^ and *rbohd* (defective for apoplastic ROS) ^36^]. We also tested wildtype Columbia-0 (Col-0) plants 6 hours after treatment with the calcium and potassium channel blocker, verapamil ^37^. Using the PCA from the RNAseq analysis (Figure 4A, Supplemental Table 1, 2), we selected four genes from the list of syringomycin- and syringopeptin-dependent genes, along with the common defense-associated gene, FRK1, which is also strongly induced in a syringomycin- and syringopeptin-dependent manner by N2C3. Furthermore, we selected five genes from the list of syringomycin- and syringopeptin-independent genes, to characterize the non-virulent defense response to N2C3. Genes were verified to have strong associations with immunity by cross-referencing with Bjornson et al., 2021 ^38^, in which genes were selected if they were induced by the majority of biotic elicitors assayed in the study, even if they were induced by some abiotic elicitors (Supplemental Table 5).

To determine if known immune pathways are required for the plant response to virulent and avirulent N2C3, in a setup similar to the RNAseq experiment, wildtype and mutant plants were treated with monocultures of virulent N2C3. Gene expression was measured in the root maturation zone two days post inoculation (DPI). The *bak1/bkk1* mutant had a significant increase in expression (p < 0.05) for the syringomycin- and syringopeptin- dependent defense-associated genes suggesting that PTI negatively regulates the response to N2C3 (Figure 5A). The *bri1* and *ein2* mutants also showed a significant increase in syringomycin- and syringopeptin-dependent reporter gene expression further suggesting that brassinosteroid signaling components and ethylene signaling may be involved in inhibiting the syringomycin and syringopeptin mediated immune response (p < 0.05). None of the other mutants showed a significant increase or decrease in reporter gene expression for the syringomycin- and syringopeptin-dependent genes. These results suggest that the syringomycin- and syringopeptin-induced defense response is not dependent on key MAMP/DAMP-PRRs, but rather is inhibited by PTI, brassinosteroid, and ethylene signaling.

When quantifying syringomycin- and syringopeptin-independent gene expression, treatment with verapamil significantly reduced reporter gene expression (p < 0.05, Figure 5F). Similar to the syringomycin- and syringopeptin- dependent genes, the *bak1/bkk1, bri1*, and *ein2*, mutants showed a significant increase in expression (p < 0.02) for the syringomycin- and syringopeptin-independent genes (Figure 5B). The remaining plant mutants did not show a significant increase or decrease in expression for the syringomycin- and syringopeptin-independent genes. These data show that the plant response to the avirulent pathogen is partially dependent on calcium or potassium signaling, and partially inhibited by bak1/bkk1, brassinosteroid, and ethylene signaling. Altogether, these results suggest that defense responses to virulent and avirulent N2C3 occur through a PTI-independent mechanism.

### Plants exhibit unique physiological immune responses to WCS365 and N2C3

Our data suggest a model in which competition between beneficial WCS365 and pathogenic N2C3 in the rhizosphere depends on complex interactions between the host and bacteria. While multiple chemical aspects of the plant rhizosphere could contribute to differences in bacterial growth, our data suggests the immune system is a likely candidate for recognizing the beneficial and pathogenic lifestyles of WCS365 and N2C3, and adjusting the plant’s physiology accordingly ^39,40^. To determine whether differences in the transcriptomic response to WCS365 and N2C3 result in unique immune responses, we use callose deposition and reactive oxygen species (ROS) burst as two physiological responses often used as outputs of immunity ^41,42^.

Prior work has shown callose to be induced via PTI ^43^. To determine if *P. fluorescens* WCS365 and N2C3 can induce callose deposition, plants were inoculated with each strain and stained with aniline blue for fluorescence-based quantification of callose deposition. Neither wildtype N2C3 nor WCS365 induced an increase in callose deposition in roots (Figure 6F). However, the *P. fluorescens* N2C3 Δ*syr*Δ*syp* mutant showed a significantly higher level of callose compared to mock in the root maturation zone (p < 0.01), suggesting a syringomycin- and syringopeptin-mediated inhibition of callose deposition.

**Figure 6.**
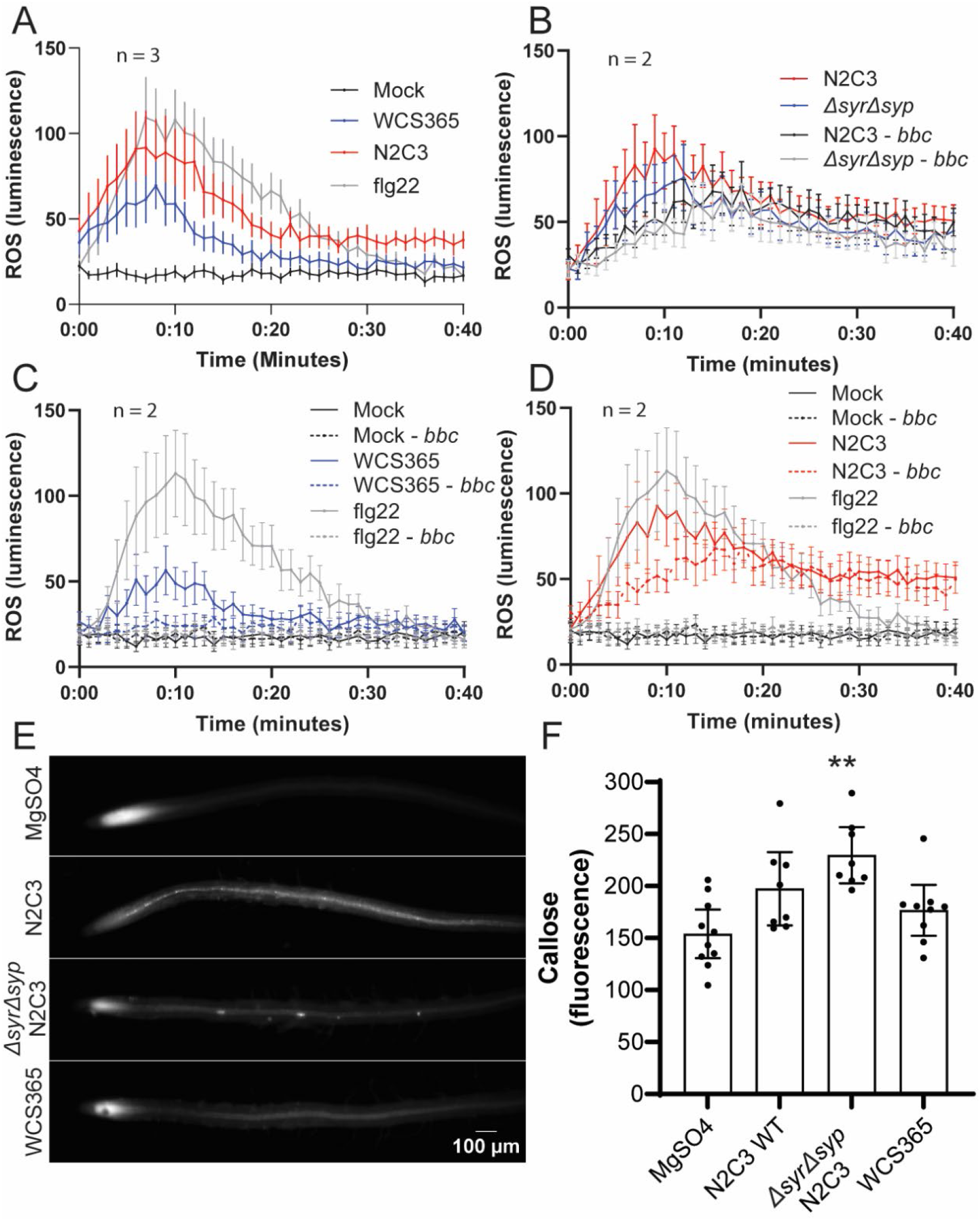
Plants exhibit a mechanistically distinct immune response to beneficial *P. fluorescens* WCS365 or pathogenic N2C3. A-D, luminescence-based assays measuring the initial ROS burst after supernatant treatment from four to six-day old WCS365 or N2C3 cultures. A, shows ROS response in wildtype plants. B, compares N2C3 wildtype and Δ*syr*Δ*syp* mutant strain induced ROS responses between Col-0 and the *bak1/bkk1/cerk1* (*bbc*) triple mutant plants. C, compares the WCS365 ROS response in Col-0 and *bbc* plants. D, compares the N2C3 ROS response in Col-0 and *bbc* plants. E and F, fluorescence-based measurement of callose deposition in the root maturation zone. E, representative fluorescence images of aniline blue stained roots two DPI. F, quantification of callose staining in the root maturation zone. N indicates the number of temporal replicates. A one-way ANOVA comparing bacterial treatments with mock was used to determine statistical significance, *p* < 0.01 = **. All error bars show 95% confidence intervals.

Prior work has shown that both commensals and pathogens can induce ROS burst via MAMP PRRs ^44^, and we found that both the beneficial *P. fluorescens* WCS365 and pathogenic N2C3 supernatant induced a rapid ROS burst in whole seedlings. The ROS burst induced by N2C3 supernatant (AUC: 2146, 95% CI: 1840 - 2451) was significantly higher than that from WCS365 supernatant (AUC: 1408, 95% CI: 1222 - 1594) (Figure 6A), and similar to that of the flg22 elicitor control (AUC: 2244, 95% CI: 1955 - 2532). This data is in agreement with our RNAseq data (Figure 3) showing that WCS365 induces a weaker defense response than N2C3.

Although both the beneficial and pathogenic strains induce immunity, our qPCR data (Figure 5) show that the transcriptomic defense response to the pathogen is not PTI dependent. Therefore, to determine whether the differences in ROS burst were due to PTI dependent and independent responses to WCS365 and N2C3, we tested the bacterial-induced ROS burst on the *bak1/bkk1/cerk1 (bbc) Arabidopsis* triple mutant which has been shown to be compromised for both PTI and ETI ^9^. We found that the WCS365 supernatant-induced ROS burst was eliminated in the *bbc* mutant (AUC: 925.3, 95% CI: 825 to 1026), exhibiting a response comparable to mock treatment (AUC: 707.7, 95% CI: 623.3 −792.0; Figure 6C). N2C3 supernatant still exhibited a strong ROS response in the *bbc* mutant (AUC: 2036, 95% CI: 1850 - 2221), which was lower than its response in Col-0 (AUC: 2428, 95% CI: 2187 - 2670) (Figure 6D). This suggests that pathogenic N2C3 elicits a potent PTI-independent immune response, accompanied by a weaker PTI mediated response. Compared to wildtype N2C3, the Δ*syr*Δ*syp* mutant supernatant induced a lower ROS burst in both Col-0 (AUC: 2031, 95% CI: 1831 - 2231) and the *bbc* mutant (AUC: 1719, 95% CI: 1548 - 1890) backgrounds (Figure 6B) indicating that the N2C3 supernatant-induced ROS is partially mediated by syringomycin and syringopeptin. Collectively, this shows that beneficial bacterial recognition is fully PTI dependent, while pathogen recognition is largely PTI independent. This indicates that plants can perceive both pathogens and beneficial strains but use distinct mechanisms to do so.

## Discussion

In this study we provide evidence that plants can distinguish between two phylogenetically similar strains of *Pseudomonas*. We show that a beneficial *P. fluorescens* induces a weak transcriptional response along with weak immune activation. A closely related *P. fluorescens* pathogen induces numerous defense-associated genes and a potent immune response. Since the beneficial strain also induces a MAMP-dependent immune response, PTI alone is not sufficient to explain the competition between the two strains in the rhizosphere. Interestingly, as PTI is often restricted to the elongation zone, the commensal-induced immunity but not pathogen-induced immunity may be spatially limited as well. In addition to bacterial MAMPs and effectors, we propose a model (Figure 7) in which immune activation can be induced by other bacterial derived virulence factors, such as the toxins presented in this study.

**Figure 7.**
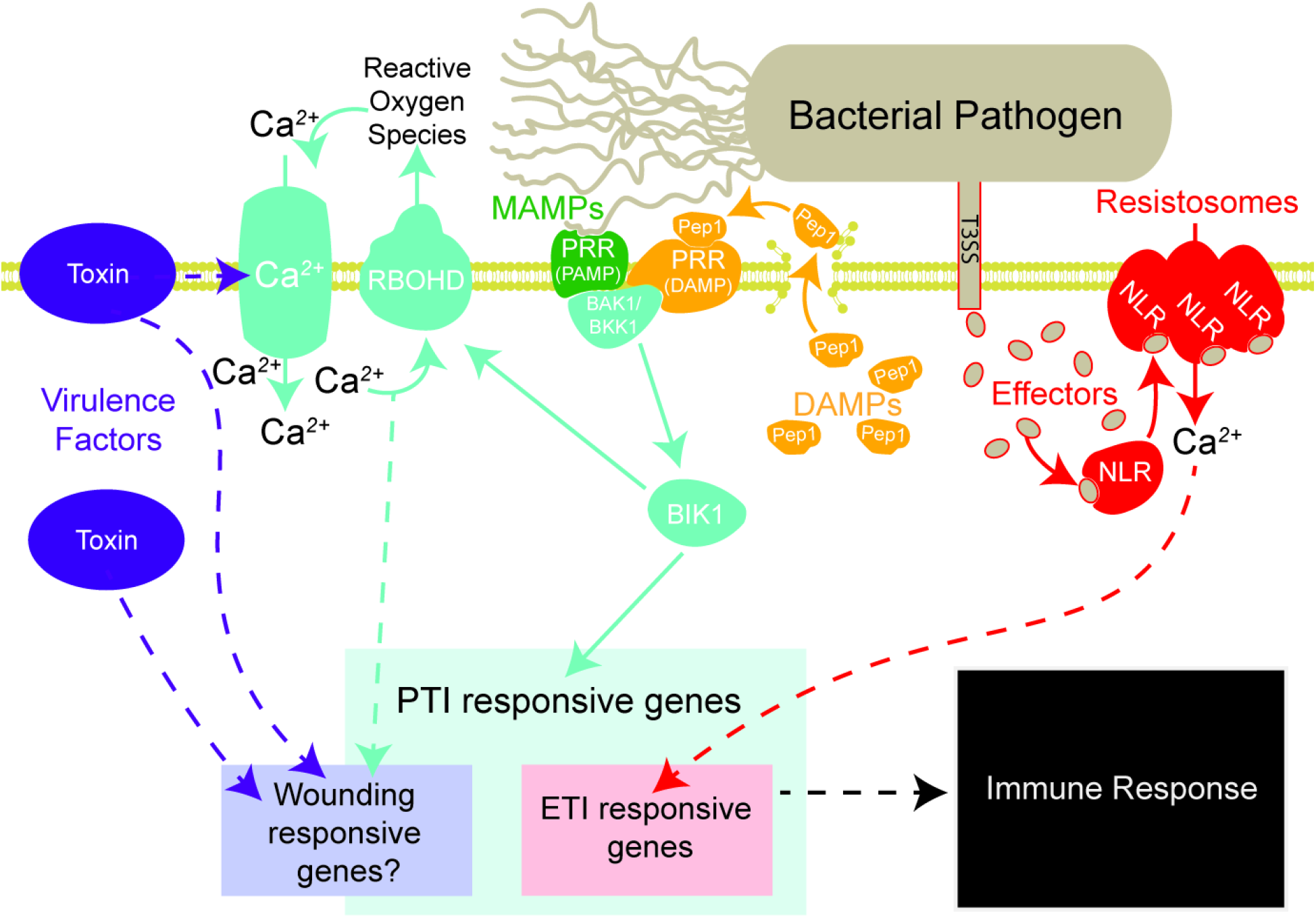
Model of how plant innate immunity may distinguish pathogens from beneficials. Plants have been shown to elicit an immune response to several types of molecules: microbe-associated molecular patterns (MAMPs), damage-associated molecular patterns (DAMPs), and effectors. Bacterial effectors are injected into the host cell via type III secretion systems (T3SS). Bacterial toxins along with other virulence factors may also elicit immunity via multiple unknown mechanisms. MAMPs induce pattern-triggered immunity (PTI) which is required for an effector-triggered immune response. Toxin-triggered immunity is not dependent on PTI but may overlap with a DAMP-induced or wounding-like defense response. However, DAMP-triggered immunity also shows considerable transcriptional overlap with PTI. Reactive oxygen species (ROS) and calcium channel influx (Ca^2+^) are key components of immunity. The activity of the toxins syringomycin and syringopeptin is dependent on Ca^2+^ influx ^64^, while resistosomes have been shown to have Ca^2+^ channel activity ^65^. Toxins may also induce ROS burst (Figure 6). Dotted arrows represent unknown pathways.

Our data suggest that pathogenic *P. fluorescens* is not inducing plant immunity via PTI or ETI, as neither the transcriptomic defense response (Figure 5) nor ROS burst (Figure 6) is PTI-dependent, and the pathogen lacks a type III secretion system necessary for inducing ETI. However, the transcriptomic response induced by the virulent and avirulent strains shares a strong overlap with AtPep1 inducible genes, suggesting the pathogen induces a damage- or wounding-like response. While important PTI components such as BAK1, BKK1, and CERK1 have recently been shown to be required for ETI ^9^, they are also required for AtPep1-triggered immunity ^45^. We find that the virulent and avirulent pathogen-triggered immune responses are not AtPep1 dependent as they are still induced in PTI deficient mutants. Prior studies show a large transcriptional overlap between immune elicitor responses ^38,46^. Therefore, while the pathogen-triggered response is not AtPep1-dependent, the large transcriptional overlap with the AtPep1 further suggests that the pathogen strongly induces immunity. Further work is needed to determine if the pathogen has other endogenous pathogenicity factors that contribute to the syringomycin-syringopeptin-independent immune response.

Our data showing a *P. fluorescens* pathogen overcomes immune barriers in the root parallels a recent study showing that PTI is restricted in the root until accompanied by a wounding-mediated signal ^24^. Interestingly, this work shows that AtPep1, among a mixture of several plant-derived damage-associated molecular patterns (DAMPs), were not sufficient to induce flg22-mediated PTI, but that wounding itself was, via an unknown mechanism. Immunity has been speculated to be induced by wounding via mechanical stimulation through sudden pressure changes in the cell ^24,47^. It should be noted that wounding is also required to initiate AtPep1-triggered immunity, by which a wound-sensing mechanism signals cleavage of AtPep1 from its precursor PROPEP1 via METACASPASE 4 ^48^. We postulate that the changes in osmotic pressure induced by bacterial virulence factors such as pore-forming toxins may also lead to similar pressure differences that parallel a wounding-mediated immune response. A wounding or damage mediated response could provide the immune system with additional information to help distinguish between harmful and beneficial bacteria.

The swift recognition of virulence factors and their effects on host physiology could allow innate immune systems to differentiate quickly evolving beneficial and pathogenic lifestyles acquired via the loss and gain of pathogenicity islands. For example, immune recognition of Shiga toxin could be useful for limiting pathogenic *E. coli* growth in the mammalian gut ^49^, while recognition of MARTX toxin effects could help control pathogenic *Vibrio sp*. growth in marine animals ^50^. Considering our finding of a PTI-independent toxin-triggered immune response, we propose that bacterial toxins may be one way by which innate immune systems may distinguish between harmful and beneficial microbes. Given bacterial toxin diversity and modes of virulence, immune recognition of toxins might occur via one or more unique mechanisms that detect perturbations in host cell physiology.

In summary, our data are consistent with the notion that plants can distinguish between harmful and beneficial microbes in a manner that is not solely reliant on evolutionarily conserved MAMPs or bacterial effectors. We propose a paradigm in which innate immune systems can distinguish between highly related pathogenic and beneficial strains in part by the presence or absence of bacterial toxins. Future work to identify genetic pathways required for these toxin-triggered responses of *P. fluorescens* N2C3, and other bacterial pathogens, will help uncover general mechanisms for how plants and animals use innate immunity to distinguish between friend and foe in microbiomes.

## Methods

### Plant growth conditions

*Arabidopsis thaliana* ecotype Columbia-0 (Col-0) is the genetic background for all wildtype, mutant, and transgenic lines. Seeds were sterilized with a 70% EtOH and 1.5% H_2_O_2_ solution on filter paper and allowed to dry in a laminar flow hood. Sterilized seeds were transferred to ½ x Murashige and Skoog (MS) square plates containing 1% agar or to a hydroponics system. All ½ x MS media contained 0.5 g/L MES buffer and was adjusted to pH 5.7. Seeds were stratified at 4°C in the dark for two to four days. Plants on agar plates were grown vertically at 22°C at 100 μM light under a 12-hour light/dark cycle.

### Hydroponics system

A hydroponics-based assay was used for *in planta* bacterial growth competitions ^51^. In brief, seeds were transferred to and germinated on floating sterile Teflon mesh suspended in 250 μL 1x MS liquid media containing 2% sucrose in clear 48-well plates. Plants were grown for 10 days before replacing the media with 270 μL ½ x MS liquid media without sucrose. 12-day old plants were inoculated with mono- or cocultures of bacteria at a final OD_600_ of 0.005.

### Bacterial strains and competition assays

Bacterial strains were transformed with modified versions of pSMC21 plasmid ^52^ for mono and co-culture assays. The plasmid pSMC21 contains 1.8kb fragment for stabilization in *Pseudomonas* sp. ^53^. The GFP gene was removed and replaced with mTurquoise2 or an XbaI restriction site to create the fluorescent reporter (pDT001) and empty vector (pDT002) plasmids, respectively (Supplemental Table 6). All bacteria were grown in LB liquid media overnight, shaking at 28°C. Cultures were washed and diluted in 10 mM MgSO_4_ prior to inoculating plants.

For each fluorescently tagged bacterial strain, linear standard curves were generated and used to estimate optical density from fluorescent intensity. Fluorescence was measure on the SpectraMax i3x plate reader, with mTurquoise2 excitation at 451 nm with a 9 nm bandwidth and emission at 486 with a 15 nm bandwidth. *In vitro* assays were performed in 96-well clear bottom plates containing ½ x MS liquid media supplemented with 30 mM succinate. Bacterial growth in both *in vitro* and *in planta* assays were measured from the bottom of the plate. *In vitro* assays were orbitally shaken for 10 seconds on medium intensity before reading at 30-minute intervals with continuous shaking between reads.

### Bacterial supernatant and elicitor treatments

Bacterial supernatants were prepared by growing bacterial cultures in a 3:1 mixture of Difco Czapek-Dox liquid media and ½ x MS liquid media. Cultures were grown at room temperature, without shaking for 3 - 6 days. Bacterial supernatant was removed after centrifugation for 10 minutes at 12,000 g at 4°C. Fresh 3:1 Cizapek-Dox:MS media was used as mock treatment.

AtPep1 (ATKVKAKQRGKEKVSSGRPGQHN) was synthesized at EZBiolab at 95% purity. Lyophilized AtPep1 was suspended in dH_2_O in 10 μM aliquots and stored at - 70°C. Flg22 was stored in 10 μM aliquots at −20°C in dH_2_O or Czapek-Dox:MS media when appropriate. For transcriptomic and microscopy experiments, plants were treated with 10 μL of 1 μM flg22 or 100 nM Atpep1 for 4.5 hours before sampling.

### Root inoculation on agar plates

Plants were grown vertically on MS agar plates for four days before all bacterial inoculations on roots grown on MS agar plates. Roots were inoculated with 2 μL of bacteria diluted to OD_600_ 0.005 and plants continued to grow vertically under a 12-hour light cycle at 23°C. Assays were performed 2 two days after inoculation.

### ROS assays

A horseradish and luminol based assay was used to measure transient ROS levels (adapted from Cheng et al. ^54^). Seeds were grown in white-bottom 96-well plates in 200 μL ½ MS media supplemented with 0.5% sucrose. After 7 days, the media was replaced with 200 μL dH_2_O. On day 8, plants were stored in the dark at room temperature for ~1 hour before treatment with bacterial supernatant or elicitors. Luminol and horseradish stock solutions were premade separately according to Cheng et al.^54^ and stored in 48 μL aliquots at −70°C. The ox-burst reagent was prepared and used under low light conditions with an aliquot of luminol and horseradish peroxidase stocks diluted into 24 mL dH_2_O at a 1:1 ratio. Plates were unwrapped, water removed from each well, and treated immediately with elicitors, supernatant, or mock. Luminescence was measured immediately on the SpectraMax i3x at 1-minute intervals and 400 ms integration times.

### RNAseq

For RNAseq sample preparation, ~30 seeds were germinated on agar ½ MS agar plates with a single plate used for each bacterial or elicitor treatment. To harvest, roots tips and shoots were quickly removed, and the remaining root tissue was immediately transferred to Eppendorf tubes containing one plastic bead. Tubes were flash frozen in liquid nitrogen and stored at −70°C until ready for RNA extraction.

Frozen tissue was ground without thawing in a TissueLyser II and prepared using a Qiagen RNeasy Plant Mini Kit according to standard protocol. Extracted RNA was DNase treated with the Invitrogen Turbo DNA-free Kit according to the standard protocol and stored at −70°C.

Processing of the Illumina sequencing data was performed with tools on usegalaxy.org using the same custom pipeline for all treatments (Supplemental Table 7), while DESeq2 was performed with R4.1.2 to identify significant (adjusted *p* value threshold of 0.01) differential gene expression.

ShinyGO GO term enrichment analysis was used to identify the enriched biological processes in the RNAseq dataset (parameters, FDR (false discovery rate) cutoff: 0.05, # pathways to show: 20, pathway size minL 5, Pathway size max: 2000, remove redundancy).

Gene set enrichment analysis (Supplemental Figure 1B, C) was done with g:Profiler, and GO terms were clustered with EnrichmentMap^55^ and annotated with AutoAnnotate^56^ in Cytoscape ^57^. CytoHubba^58^ was used to identify important biological nodes within the “Defense Response” biological process GO term using the MCC method.

### Callose deposition assay

The fixative was prepared with a 3:1 mix of 95% ethanol to acetic acid (adapted from Amherst et al., 1996; Eschrich and Currier, 2009). The fixative was cooled to approximately −74°C by slowly adding crushed dry ice. A 6-well plate was cooled on dry ice and liquid nitrogen was added to each well. After flash freezing two-day post inoculated plants in the appropriate wells, 3 mL of the fixative was added. Plates were kept on dry ice and stored at −20°C for 2-8 hours before transferring to 4°C overnight. Plates were then slowly brought to room temperature over a 1–2-hour period. Plants were washed several times over a 1.5 – 2-hour period while shaking at 120 rpm. Aniline blue was prepared in K_2_HPO_4_, and samples were stained in 0.005% aniline for 30 minutes. Samples were mounted in 50% glycerol in K_2_HPO_4_ and imaged on a MVX10 fluorescence stereomicroscope.

### RT-qPCR setup

RNA for qPCR was prepared in the same manner as for RNAseq. The Invitrogen Qubit was used to quantify RNA. The Invitrogen SuperScript III First-Strand Synthesis System (Catalog #18080051) was used according to standard protocol to prepare cDNA. SYBR green was used for RT-qPCR. Each 10 μL reaction used a final concentration of 2.5 ng cDNA, 400 nM forward and reverse primers (Supplemental Table 5), 1x PowerUp SYBR Green Master Mix (A25742), and dH_2_O.

RT-qPCR reactions (Figure 5) were prepared using the Echo acoustic dispenser in 384-well plates. RT-qPCR was run on the QuantStudio 6 Pro using a Standard run mode and relative quantification. Initial denaturation was at 95°C for 2 minutes. Denaturation was at 95°C for 15 seconds and annealing and extension was at 60°C for 1 minute for 40x cycles.

### Confocal Microscopy

The Leica SP5 laser scanning confocal was used for imaging and quantification of the pPER5::mVenusx3-NLS fluorescence reporter. Plants were mounted on 1% agar in a modified glass slide imaging chamber ^61^ and covered with a VWR 22×50 mm #1.5 glass coverslip. 512×512 pixel, 8-bit images were acquired with a 60x 1.4 NA oil objective. The Argon laser power was set at 70% and the 514-laser line was set to 22%. PMT detectors captured fluorescence at 524 – 648 nm bandwidth. The pinhole was set to 2.5 Airy Units (238.8 μm), and optically sectioned at 1.73 μm step-size. Fluorescence images were quantified with FIJI is Just ImageJ. Images stacks were summed, and the segmented line tool (width adjusted for root width) was used to trace the length of a root and the Plot Profile tool was used to measure fluorescence.

## Acknowledgements

RNA sequencing and library preparation was performed with the expertise of Malgorzata Michelle Moksa. Setup of RT-qPCR experiments via the Echo acoustic dispenser was performed by Nicholas Ek. Microscopy was performed at University of British Columbia Bioimaging Facility and LSI Imaging core. We thank Dr. Niko Geldner for the mVenus plant immune reporter lines and Dr. Yuelin Zhang and the Dr. Cyril Zipfel lab for the *bak1/bkk1* and *bbc* triple mutant lines. We thank Daniela Yanez for discussions on genomic analysis tools. This work was supported by the NSF National Plant Genome Initiative Postdoctoral Research Fellowship in Biology (#2010946) awarded to D.T., an NSERC Discovery Grant and Accelerator Award (NSERC-RGPIN-2021-03587) and a Canada Research Chair salary award to C.H.H. Additional trainee support was provided by Chinese Scholarship Council Awards to Y.L. and S.S., and an NSERC CGS-M award to N.R.W.

## Supplemental Figure

**Supplemental Figure 1.**
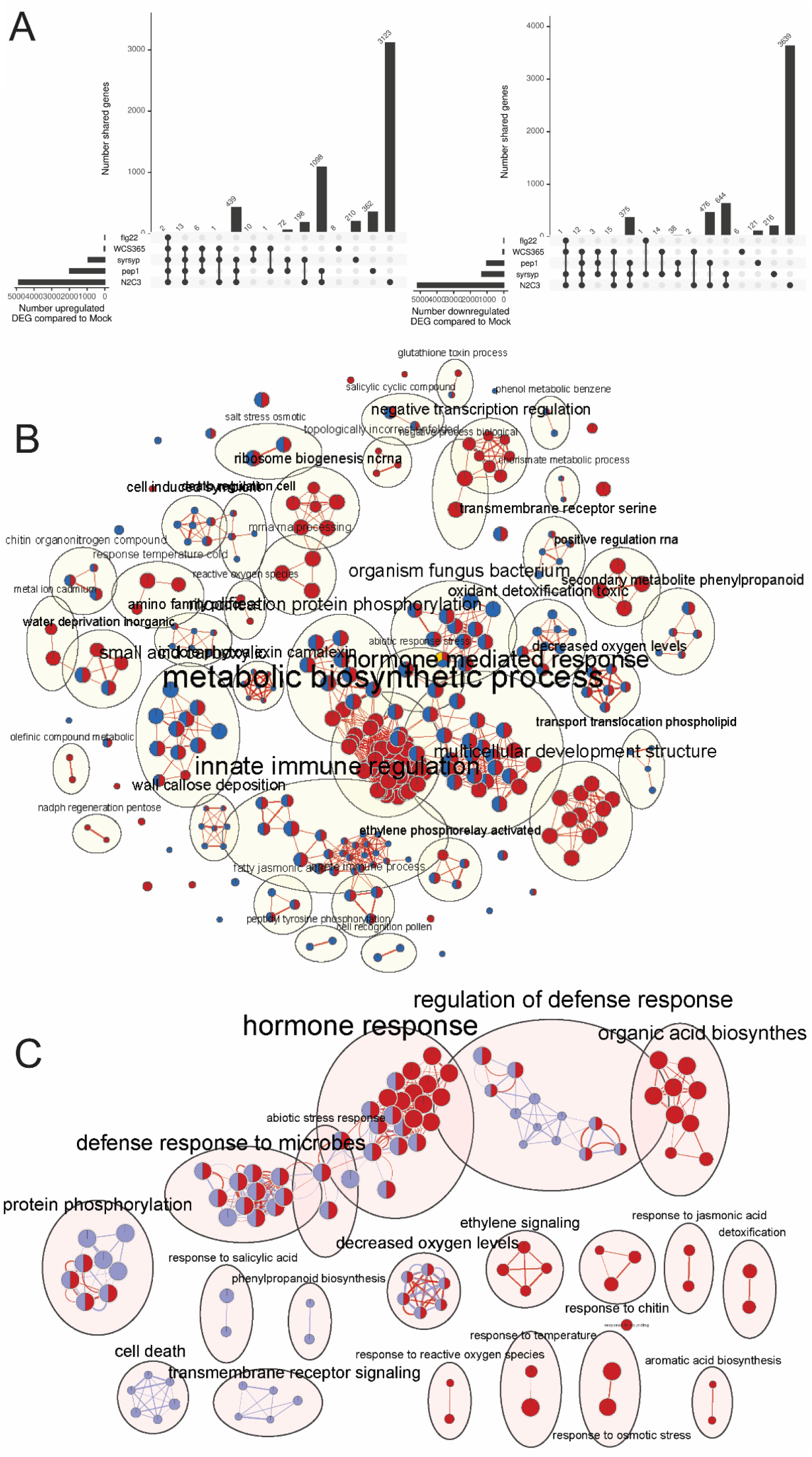
A, Upset plots showing the number of upregulated or downregulated differentially expressed genes (DEGs) in the root maturation zone comparing treatments with WCS365, N2C3, Δ*syr*Δ*syp* N2C3, flg22, or AtPep1, to mock. B, Gene set enrichment analysis of significant DEGs from the total RNAseq dataset (Figure 3A) showing the overlap between N2C3 (red nodes), AtPep1 (blue nodes), and WCS365 upregulated DEGs (yellow node). C, Gene set enrichment analysis of the PCA data (Figure 4A) showing the overlap between wildtype N2C3 (red nodes) and ΔSYRSYP N2C3 mutant (purple nodes). Significant DEGs were selected based on the DESeq2 adjusted *p* value (*p* < 0.01).

